# A pseudotyped lentivirus-based assay to titer SARS-CoV-2 neutralizing antibodies in Mexico

**DOI:** 10.1101/2022.01.27.478128

**Authors:** José Antonio Cruz-Cardenas, Michelle Gutierrez-Mayret, Alejandra López-Arredondo, Julio Enrique Castañeda-Delgado, Augusto Rojas-Martinez, Gerardo García-Rivas, José Antonio Enciso-Moreno, Laura A. Palomares, Marion E. G. Brunck

## Abstract

Measuring the neutralizing potential of SARS-CoV-2 antigens-exposed sera informs on effective humoral immunity. This is relevant to 1-monitor levels of protection within an asymptomatic population, 2-evaluate the efficacy of existing and novel vaccines against emerging variants, 3-test prospective therapeutic monoclonal neutralizing antibodies (NAbs) and, overall, to contribute to understand SARS-CoV-2 immunity. However, the gold-standard method to titer NAbs is a functional assay of virus-mediated infection, which requires biosafety level 3 (BSL-3) facilities. As these facilities are insufficient in Latin American countries, including Mexico, scant information has been obtained about NAb in these countries during the COVID-19 pandemic. An alternative solution to acquire NAb information locally is to use non-replicative viral particles that display the SARS-CoV-2 Spike (S) protein on their surface, and deliver a reporter gene into target cells upon transduction. Here we present the development of a NAb-measuring assay based on Nanoluc-mediated luminescence measurements from SARS-CoV-2 S-pseudotyped lentiviral particle-infected cells. The successive steps of development are presented, including lentiviral particles production, target cell selection, and TCID50 determination. We applied the optimized assay in a BSL-2 facility to measure NAbs in 15 pre-pandemic, 18 COVID-19 convalescent and 32 BNT162b2 vaccinated serum samples, which evidenced the assay with 100% sensitivity, 86.6% specificity and 96% accuracy. The assay highlighted heterogeneity in neutralization curves which are relevant in discussing neutralization potency dynamics. Overall, this is the first report of a BSL-2 safe functional assay to measure SARS-CoV-2 in Mexico and a cornerstone methodology necessary to measure NAb with a functional assay in the context of limited resources settings.

**Importance:** Evaluating effective humoral immunity against SARS-CoV-2 requires a functional assay with infectious virus. Handling the authentic SARS-CoV-2 virus requires specialized facilities that are not readily available in Latin America, including Mexico. Here we produce non-replicative viral particles pseudotyped with the SARS-CoV-2 S protein that are used as safe surrogate viral particles in an optimized BSL-2 ready neutralization assay. The establishment of this assay is critical to allow the evaluation of effective humoral immunity to SARS-CoV-2 post-infection and to monitor the efficacy of existing or novel vaccines against emerging variants in the Mexican population.

## Introduction

Humoral immunity provides critical protection against viruses including memory against future infections. In particular, neutralizing antibodies (NAbs) specifically target epitopes on viral membrane proteins, interfering with cell receptor binding^1^. NAbs against SARS-CoV-2, the causal agent for COVID-19, interfere with viral infection by various potential mechanisms, all culminating in preventing viral entry^2^. One of these mechanisms is the binding of the receptor binding domain (RBD) of the S1 subunit of the Spike protein (S) of SARS-CoV-2, which hampers interactions with the angiotensin converting enzyme 2 (ACE2) receptor on target cells, and therefore blocks viral entry^3^.

During the natural course of SARS-CoV-2 infection, the early emergence of NAbs prevents fatal disease^4^. Vaccine induced SARS-CoV-2 NAbs correlate with increased survival, decreased symptoms severity and reduced risk of re-infection^5,6^. In addition, monoclonal NAbs prevent SARS-CoV-2 infections *in vitro* and *in vivo*. Various monoclonal NAbs have been given emergency use authorization by the FDA, such as casirivimab and imdevimab, while others and are the subject of current clinical trials as therapeutic prospects^7–10^. However, the regular emergence of variants of concern such as B.1.617.2 (Delta) and more recently B.1.1.529 (Omicron), motivates continuing vaccine- and directed therapeutic-research efforts worldwide, including in Mexico^11–13^.

To monitor the development and prevalence of an effective humoral response against SARS-CoV-2 and emerging variants in a population, it is therefore essential to measure SARS-CoV-2-specific NAbs, generated either through natural infection, or through the application of vaccines. Measuring NAbs requires a functional assay whereby serum samples are co-incubated with SARS-CoV-2 viral particles (VP), after which the ability of these VP to infect target cell *in vitro* is measured^14^.

The use of SARS-CoV-2 reference strains or clinical isolates for NAb titration experiments requires Biosafety Level 3 (BSL-3) laboratories, which are scarce in Latin America. As a result, few reports are available about SARS-CoV-2 NAb in the Mexican population, with all currently available data relying on a neutralization-surrogate ELISA kit^15–17^. During a pandemic, access to specific research reagents, such as an imported ELISA kit, is limited which cause long delays to the evaluation of the effectiveness of the national vaccination program and to the monitoring of the seroprevalence of the disease, both important aspects in the control of COVID-19.

An alternative strategy to using authentic virus is to produce non-replicative VP that express the SARS-CoV-2 S or RBD on their surface and that include a reporter gene delivered to target cells upon transduction^18–20^. These pseudotyped VP have been widely used to measure NAbs against a range of potentially fatal viruses, including influenza (H7N9), MERS-CoV, HCV, and SARS-CoV-2 and recent variants^21–24^. Advantages of using pseudotyped VP to measure NAbs include the facility of upscaling for high-throughput measurements at a lower cost, as well as the opportunity to customize the viral glycoprotein to match emerging variants.

Various viral backbones have been used to produce VP that express SARS-CoV-2 S in their membrane, including rhabdoviruses (VSV), retroviruses (MLV), and lentiviruses (HIV-1)^18,19,25^. Genomes of pseudotyped VP are modified to prevent viral replication, and include reporter genes such as GFP or luciferases (like fLuc or Nluc), that facilitate transduction monitoring in target cells^14,26^. Lentiviral systems constitute a popular backbone to manufacture pseudotyped VP due to their short production time, relative high yields and ease of handling^27^. Lentiviral systems are available in 2^nd^, 3^rd^ and 4^th^ generation systems that allow their safe manipulation in BSL-2 laboratories^28^.

Here we present the development of a SARS-CoV-2 NAb titration assay based on non-replicative pseudotyped lentiviral particles integrating Nluc into transduced cells genomes. The assay facilitated quantification of effective humoral immunity to SARS-CoV-2 in COVID-19 convalescent patients and BNT162b2 vaccinated individuals. The assay could easily be deployed in BSL-2 laboratories to investigate humoral immunity in infected patients, effective protection from vaccine administration, and to support therapeutics and vaccine development and research efforts locally.

## Results

### Development and production of a SARS-CoV-2 pseudotyped lentivirus

To develop a BSL-2-ready assay to investigate neutralizing antibodies to SARS-CoV-2 in Mexico, we first produced SARS-CoV-2 S-pseudotyped VP. We optimized a previously reported 3^rd^-generation lentiviral system (Fig. 1) by using the reporter gene Nluc which is more stable and provides 100x brighter luminescence compared to fLuc^17,20^. SARS-CoV-2 pseudotyped lentiviral vectors were achieved by incorporating a S sequence that lacks the last 19 amino acids at the C-terminal, reported to increase its incorporation into pseudoviral membranes compared to the original sequence (Fig. 1A)^25^. The Nluc gene was cloned into the transfer plasmid within LTRs to allow efficient integration in target cells upon viral entry (Fig. 1C). Additional plasmids were produced as controls, to express either the glycoprotein of VSV virus (VSV-G) that binds to the ubiquitously expressed low-density lipoprotein (LDL), or no glycoprotein (Fig. 1A) ^18,23^. The integrity of all the constructions used in this work was verified by Sanger sequencing with 100% identity (Fig. S1).

**Fig. 1.**
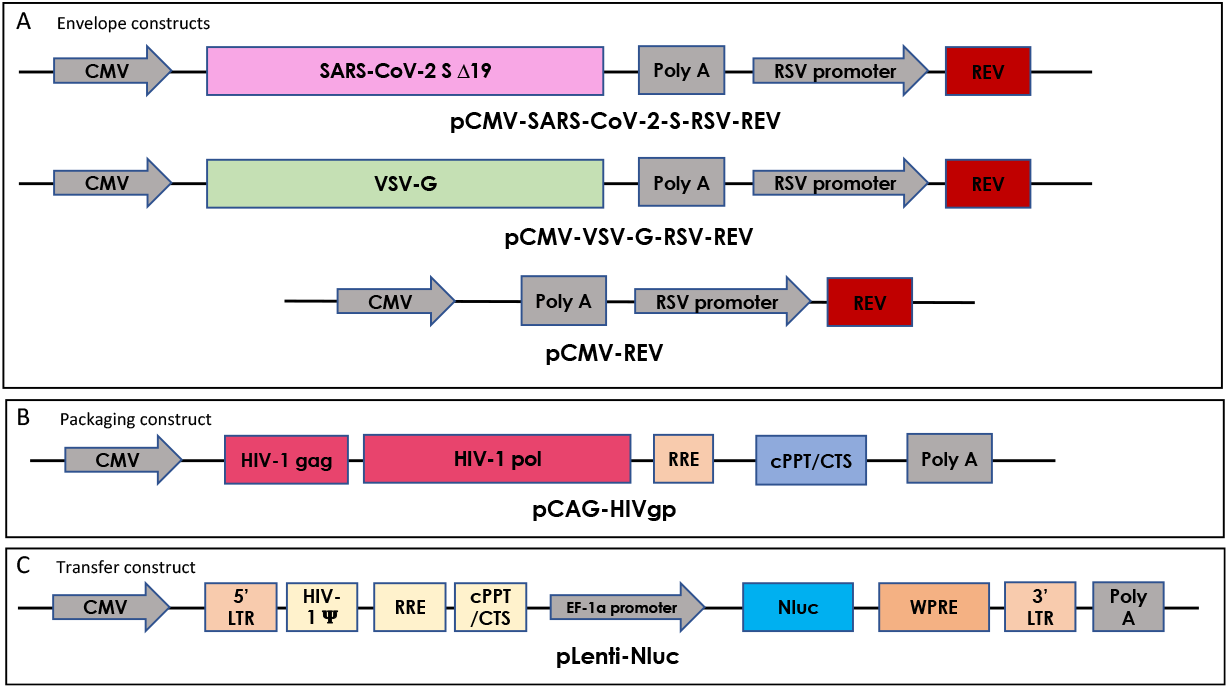
Schematic representation of constructs developed as part of a third-generation lentiviral-based system to produce VP pseudotyped either with SARS-CoV-2 S, VSV-G or no envelope protein.

The selective expression of relevant viral proteins in produced VP was investigated using western blots. The presence of the structural protein p24, core component of the lentiviral particles, was confirmed in the 3 types of VP produced (Fig. 2A)^24^. A specific band of 70 kDa was observed selectively in the VSV-G VP sample, which is consistent with the expected size of VSV-G (Fig. 2B)^29^. The selective incorporation of the S protein in SARS-CoV-2 S VP was confirmed with the detection of a 110 kDa band, using a chimeric monoclonal antibody, while in the same gel, no protein was detected for the VSV-G expressing VP (Fig. 2C)^20^.

**Fig. 2.**
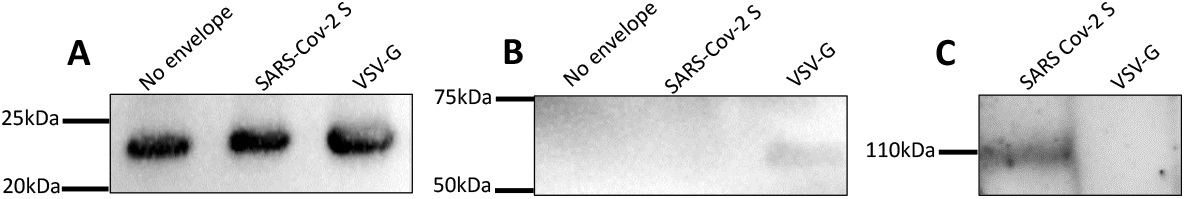
Western blot analysis of produced VP. A: The structural protein p24 was detected on the 3 types of VP. B: VSV-G was selectively detected on VSV-G VP but not on either SARS-CoV-2 S or no-envelope protein VP. C: The S protein was detected as a 110 kDa protein on SARS-CoV-2 VP using a chimeric monoclonal antibody, but not on VSV-G VP. Detection of viral proteins was performed twice and one is shown.

### Optimization of a SARS-CoV-2 pseudovirus-based neutralization assay

ACE2 expression on cell surfaces correlates with SARS-CoV-2 infection susceptibility *in vitro*^30^. Therefore, we sought to select the most appropriate target cell line for the infection assay by investigating ACE2 expression on exposed cell membranes of various cell lines known to endogenously express ACE2 and that have been previously reported as target cells for SARS-CoV-2 and pseudotyped VP^23,31–33^. Caco-2 showed 16% of ACE2 positivity in culture, while Vero and Vero E6 were homogeneously highly expressing cells with 92% and 73% positivity, respectively (Fig. 3A). Heterogeneity in ACE2 expression was recently reported between single cells of various cell lines, with expression being modulated during culture and regulated epigenetically^34^. Here, Vero cells had a significantly higher expression of ACE2 on cell surfaces compared to all other cell lines tested while HEK-293T lacked expression of ACE2 (Fig. 3B), as previously described^35^. Surface expression of ACE2 suggested that Vero cells would be the most susceptible to infection by SARS-CoV-2 pseudotyped-VP. Other techniques have been applied with mixed results to investigate ACE2 expression by target cells, including Western blot and qRT-PCR, both precluding distinction between membrane-displayed and cytosolic stores of ACE2 in contrast with flow cytometry^36,37^. However, as additional membrane proteins such as TMPRSS2 and neuropilin-1 have been evidenced as SARS-CoV-2 VP entry facilitators, the expression of ACE2 may not be sufficient on its own to predict susceptibility to infection^19,34,38^.

**Fig. 3.**
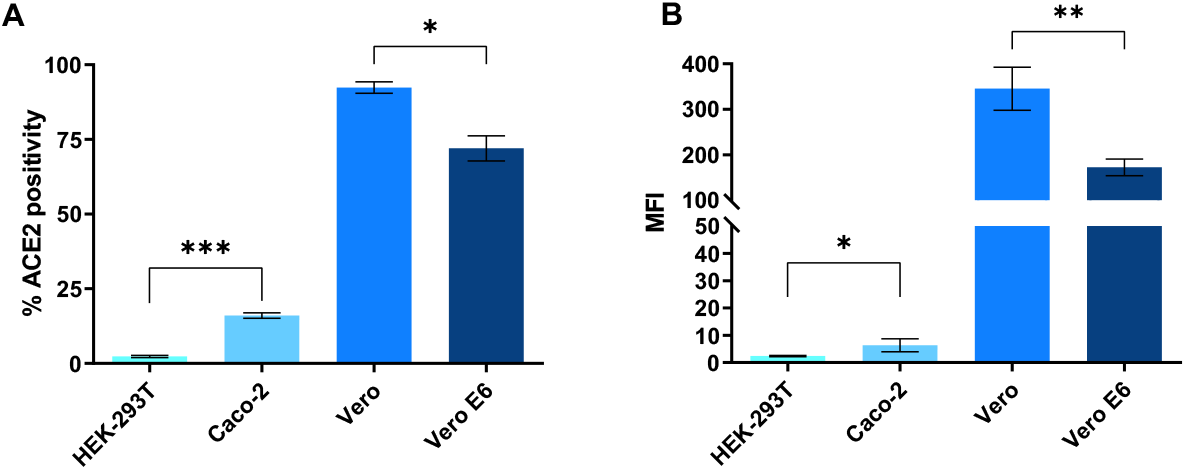
Expression of ACE2 on the surface of HEK-293T, Caco-2, Vero, and Vero E6 cells. A: Proportion of cells expressing ACE2 B: Relative expression of ACE2 on cell surface as expressed by median fluorescence intensity (MFI) of ACE2-AF647. Average of 3 separate experiments done in duplicate with standard deviations shown. p <0.05 (*), p <0.01 (**), p <0.001 (***).

Accordingly, to compare infection susceptibility of ACE2-expressing cell lines by SARS-CoV-2 pseudotyped-VP, we co-cultured Caco-2, Vero and Vero E6 with SARS-CoV-2 pseudotyped-VP and measured Nluc activity as a surrogate marker for infection. Vero cells produced the highest RLUs (mean = 9.5 x10^5^, 8.4 x10^5^ and 6.8 x10^5^ RLUs for Vero, Vero E6 and Caco-2, respectively, Fig. 4A), consistent with high ACE2 expression on cell surfaces (Fig. 3). Vero and Vero E6 exhibited similar susceptibility to VSV-G with an average of 9 x10^5^ and 8.7 x10^5^ RLUs, respectively. On the other hand, Caco-2 showed a third of the Vero lines response (mean = 3.1 x10^5^ RLUs, Fig. 4B). Interestingly, a previous report using a VSV-pseudotyped VP system showed Caco-2 had similar susceptibility to VSV-G VP and SARS-CoV-2 VP^19^. However, the complete SARS-CoV-2 S sequence was used in this case, while in the present work the use of a SARS-CoV-2-S Δ19 sequence which increases the incorporation of S in VP membranes, statistically augments opportunities for ACE2 binding and cell entry, and increasing the susceptibility. This is a possible cause for the reported differences in VSV-G-mediated and SARS-CoV-2-S Δ19-mediated infections of Caco-2 cells. Culturing cell lines together with VP that lacked surface glycoprotein led to a 30-fold decrease in infection rates, compared to infections with SARS-CoV-2 S VP, with RLUs consistently < 3 x10^4^, as expected^3^. Observed basal levels of non-specific viral entry have been reported and may be a consequence of endocytosis^35^. Overall, we replicated previous findings confirming Vero cells are highly susceptible to transduction with SARS-CoV-2 VP, leading further experiments to be performed with the Vero cell line^24^. To identify the amount of VP required to transduce 50% of the culture (TCID50), as relevant to investigate the effect of an inhibitor in biological assays, we performed a serial dilution of the SARS-CoV-2 pseudotyped-VP and applied the Reed-Muench method (Sup. Fig. 2)^39,40^. We identified 15 pg SARS-CoV-2 pseudotyped-VP were necessary to infect 50% of 25,000 Vero cells in a 96-well plate at 24 h post-inoculation.

**Fig. 4.**
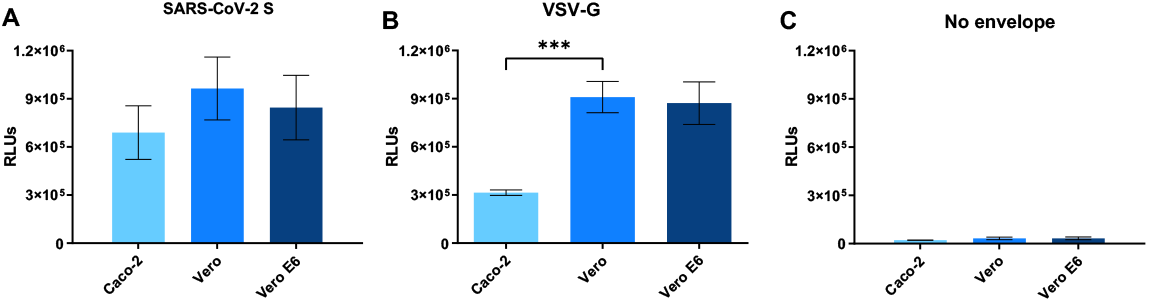
Transduction levels in Caco-2, Vero, and Vero E6 cells 24 h post-inoculation with VP, as evidenced by luminescence in relative light units (RLUs) caused by Nluc digestion of a furimazine substrate. The 3 cell lines were infected with 140 pg VP pseudotyped using A: SARS-CoV-2 S, B: VSV-G, or C: no envelope glycoprotein. Results represent the average of 3 separate experiments performed in duplicate, and standard deviations of these results are shown, p <0.001 (***).

### Neutralization of SARS-CoV-2 S lentiVP by convalescent and vaccinated sera

Once assay parameters were optimized, neutralization of transduction with human sera was implemented. Sera collected prior to the start of the COVID-19 pandemic, sera from COVID-19 diagnosed patients, and sera from health professionals that had received the BNT162b2 vaccine were used (Tables 1 and 2). Pre-pandemic sera showed neutralization of the SARS-CoV-2 pseudotyped-VP ranging between 11.6% and 41% at the lowest (1:5) dilution tested, and ranging between 20.2% and 4.6% at the highest tested dilution (1:9860, Fig. 5A). These results are consistent with the literature, where dilution-dependent, consistent, minimal cross-neutralization of SARS-CoV-2 VP by pre-pandemic sera are reported but considered insignificant for preventing COVID-19^18,23,41^. In another report, antibodies produced by various B cell clones obtained from a SARS-CoV 2003 outbreak survivor could efficiently neutralize SARS-CoV-2 and a SARS-related bat virus suggesting some levels of cross-neutralization^42^. All sera with prior exposure to SARS-CoV-2 (through natural infection or vaccination) could neutralize SARS-CoV-2 pseudotyped-VP efficiently. Convalescent COVID-19 sera were heterogeneous in their neutralization potential, with the lowest dilution (1:5) neutralizing between 95.9% and 58.5% of infection, and half of tested serum samples had a 30% neutralization titer of 540 (Fig. 5B, Table1). Of note, the similar neutralization rates observed at the 1:5 and 1:20 dilutions (means = 76.25 % and 74.7%, SD = 13. 6 and 11.6, respectively) suggest that overall NAbs contained in COVID-19 diluted down to 1:20 were in excess over VP. In contrast with convalescent sera, 14 out of 16 vaccinated samples (87.5%) had a 30% neutralization titer of 540 after the first BNT162b2 dose, and all samples had a 30% neutralization titer >540 post-boost. Six out of the 16 individuals in the vaccinated cohort had COVID-19 positive diagnostic prior to vaccination without notable impact on reported neutralization rates (Table 2). Two individuals showed slight decrease in neutralization after the second dose (Table 2), which has been reported before^16^. Importantly, the potency of vaccinated sera was higher than COVID-19 sera, with >18% vaccinated sera (3/16) having a 30% neutralization titer of 9860, versus only 5.5% COVID-19 sera (1/18). Using these values the presented assay has a 100% sensitivity, 86.6% specificity and 95.9% accuracy (Fig. 5D) using 1:20 serum dilution, as previously reported for such calculations ^43^. Looking at individual neutralization curves, there was no consensus pattern across serum dilution (Fig. 6). Some samples exhibited similar % neutralization at the lowest dilution after the first and 2^nd^ vaccine doses (as shown in representative samples ID 43, ID 46), while others evidenced increased neutralization at the lowest dilution after the 2^nd^ dose (exemplified by ID 44, and to a lower extent ID 42). The shapes of neutralization curves could be concave with a slow decrease in neutralization before reaching EC50 (ID 39, ID 42, ID 21), or convex with a sharp slope around EC50 (ID 41, ID 44) and these differences could be observed within a same individual, between 2 samples (ID 43). Interestingly, the curve of convalescent patient ID 21 exhibited constant, moderately high neutralization >75% between 1:5 and 1: 540 serum dilutions, followed by a sharp decrease in neutralization between serum dilutions 1:540 and 1:9860. Due to the shape of this curve, EC50 is extremely low suggesting very potent neutralization serum, however the patient suffered severe symptoms and passed away (Table 1).

**Table 1:** Clinical parameters and neutralization information from COVID-19 convalescent sera

| ID | COVID-19<br>diagnostic<br>q-PCR validated | Gender | Age | Days after initiation<br>of symptoms<br>at the time of sampling | Severity | Hospitalization status | Artificial<br>respiration | Patient<br>outcome | EC50 | R<sup>2</sup> | % Neutralization<br>1:20 sample dilution |
| --- | --- | --- | --- | --- | --- | --- | --- | --- | --- | --- | --- |
| 2 | Positive | M | 42 | 0 | Asx/Mild | Outpatient | NA | Recovered | 0.001401 | 0.972 | 83.03 |
| 4 | Positive | F | 65 | 19 | Asx/Mild | Outpatient | NA | Recovered | 0.004868 | 0.977 | 69.01 |
| 7 | Positive | F | 76 | 17 | Asx/Mild | NA | NA | Recovered | 0.023690 | 0.959 | 66.17 |
| 10 | Positive | M | 58 | 11 | Asx/Mild | Outpatient | NA | Recovered | 0.004090 | 0.861 | 70.25 |
| 11 | Positive | M | 58 | 30 | Asx/Mild | Outpatient | NA | Recovered | 0.006593 | 0.974 | 80.66 |
| 15 | Positive | F | 62 | 26 | Severe | Severe | NA | Recovered | 0.007370 | 0.996 | 67.46 |
| 18 | Positive | M | 53 | 77 | Severe | Severe | NA | Recovered | 0.011600 | 0.998 | 78.39 |
| 21 | Positive | M | 42 | 16 | Severe | Severe | NA | Deceased | 0.000000031 | 0.976 | 76.02 |
| 23 | Positive | M | 43 | 18 | Severe | Severe | NA | Recovered | 0.018150 | 0.998 | 83.03 |
| 25 | Positive | M | 59 | 19 | Severe | Severe | NA | Deceased | 0.013700 | 0.966 | 58.58 |
| 27 | Positive | M | 36 | 9 | Severe | Severe | Yes | Deceased | 0.003756 | 0.998 | 59.37 |
| 28 | Positive | M | 36 | 16 | Severe | Severe | Yes | Deceased | 0.005466 | 0.935 | 71.41 |
| 29 | Positive | M | 36 | 23 | Severe | Severe | NA | Deceased | 0.000934 | 0.976 | 95.92 |
| 31 | Positive | M | 34 | 19 | Asx/Mild | Outpatient | NA | Recovered | 0.009537 | 0.944 | 86.85 |
| 32 | Positive | F | 35 | 21 | Asx/Mild | Outpatient | NA | Recovered | 0.001044 | 0.996 | 89.45 |
| 33 | Positive | M | 66 | 17 | Severe | Severe | NA | Recovered | 0.010410 | 0.990 | 85.83 |
| 34 | Positive | M | 52 | 21 | Severe | Severe | NA | Recovered | 0.002027 | 0.979 | 92.59 |
| 36 | Positive | M | 49 | 9 | Asx/Mild | Outpatient | NA | Deceased | 0.010780 | 0.983 | 77.48 |

NA: Unavailable information ; Asx : asymptomatic. Highlighted cells represent longitudinal samples from same patients (patient male 58 y.o, sample IDs 10-11; Patient male 36 y.o, sample IDs 27-29).

**Table 2:** Clinical and neutralization information from BNT162b2 vaccinated individuals

| ID | COVID-19<br>Diagnosis | Gender | First dose |  |  |  | Boost |  |  |  |
| --- | --- | --- | --- | --- | --- | --- | --- | --- | --- | --- |
|  |  |  | Days post vaccine<br>administration<br>at the time of sampling | EC50 | R<sup>2</sup> | % Neutralization<br>1:20 dilution | Days post vaccine<br>administration<br>at the time of sampling | EC50 | R<sup>2</sup> | % Neutralization<br>1:20 dilution |
| 39 | Negative | F | 16 | 0.001683 | 0.992 | 90.14 | 28 | 0.000554 | 0.986 | 96.93 |
| 40 | Negative | F | 14 | 0.002394 | 0.999 | 92.78 | 29 | 0.002260 | 1.000 | 96.44 |
| 41 | Negative | F | 14 | 0.005097 | 0.990 | 88.19 | 29 | 0.003490 | 0.996 | 84.59 |
| 42 | Positive<br>recovered | M | 14 | 0.001225 | 0.996 | 89.29 | 28 | 0.000724 | 0.999 | 96.60 |
| 43 | Negative | M | 14 | 0.005883 | 0.988 | 92.08 | 29 | 0.000278 | 1.000 | 97.21 |
| 44 | Negative | F | 16 | 0.004726 | 0.989 | 67.14 | 28 | 0.003606 | 1.000 | 90.52 |
| 45 | Negative | M | 16 | 0.001136 | 0.995 | 93.80 | 28 | 0.002407 | 0.992 | 97.02 |
| 46 | Negative | F | 16 | 0.000439 | 0.994 | 93.57 | 27 | 0.001298 | 0.999 | 96.91 |
| 47 | Positive<br>recovered | M | 16 | 0.000781 | 0.951 | 93.25 | 27 | 0.000683 | 0.993 | 95.54 |
| 48 | Negative | F | 16 | 0.000916 | 0.979 | 98.04 | 27 | 0.000940 | 1.000 | 97.59 |
| 49 | Negative | F | 14 | 0.001746 | 0.995 | 97.60 | 28 | 0.001864 | 0.992 | 97.51 |
| 50 | Negative | F | 14 | 0.001030 | 0.986 | 97.36 | 28 | 0.001096 | 0.999 | 98.07 |
| 51 | Positive<br>recovered | F | 14 | 0.003675 | 0.980 | 92.60 | 28 | 0.001998 | 0.997 | 97.16 |
| 52 | Positive<br>recovered | F | 13 | 0.001203 | 0.998 | 96.58 | 27 | 0.004783 | 0.997 | 78.76 |
| 53 | Positive<br>recovered | F | 16 | 0.000000 | 0.998 | 91.51 | NA | 0.000967 | 0.975 | 97.56 |
| 54 | Positive<br>recovered | F | 16 | 0.003309 | 0.981 | 89.43 | 28 | 0.001489 | 0.981 | 95.23 |

NA: Unavailable information

**Fig. 5:**
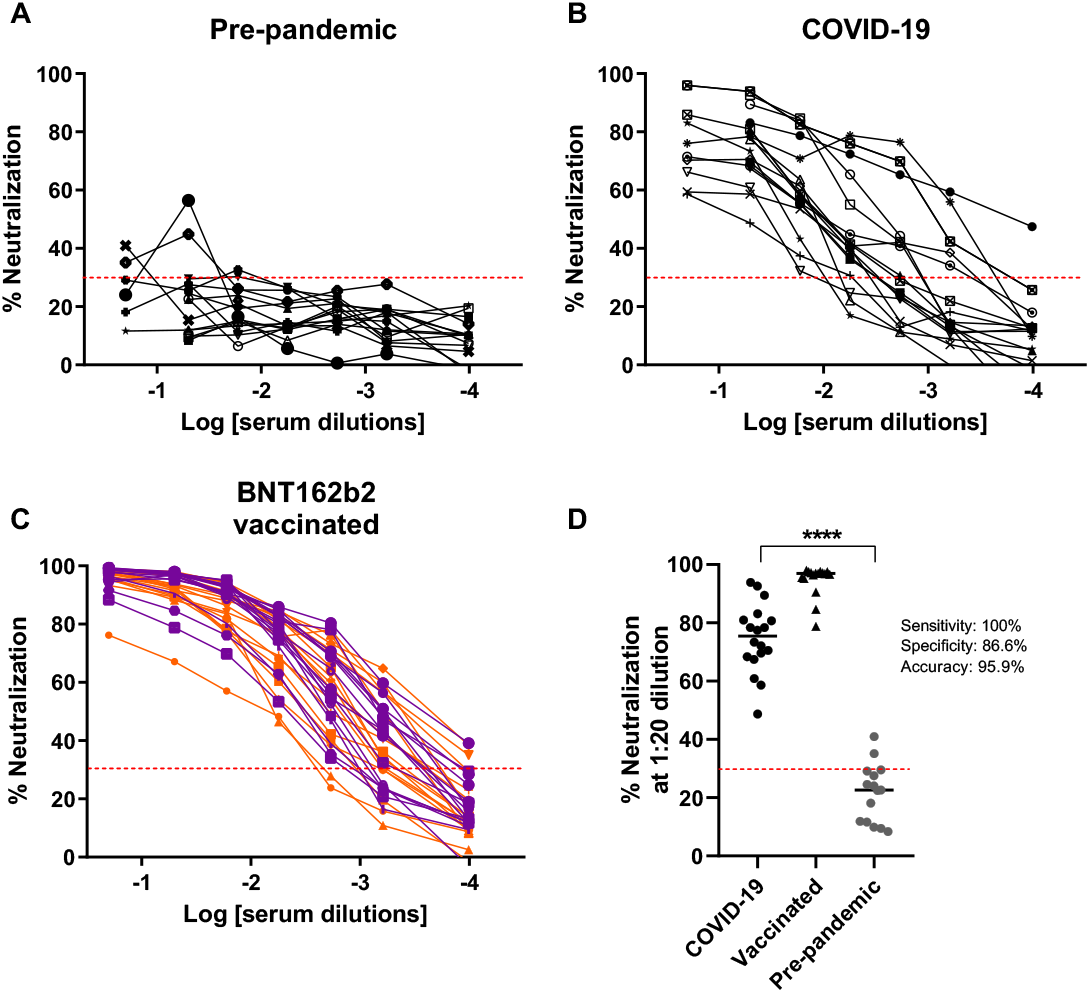
Measurement of SARS-CoV-2 S pseudotyped VP-neutralization activity from human samples. A: Pre-pandemic sera. B: Covid-19 convalescent sera. C: BNT162b2 vaccinated sera. Orange: sample obtained on average 19 days [range: 13-22] after the first vaccine dose, purple: sample obtained days on average 26 days [range: 22-29] after the 2^nd^ vaccine dose. D: Calculations of assay sensitivity, specificity and accuracy using neutralization results at 1:20 sera dilutions. The dotted line represents an arbitrary 30% neutralization cutoff for titers estimation. p <0.0001 (****).

**Fig. 6:**
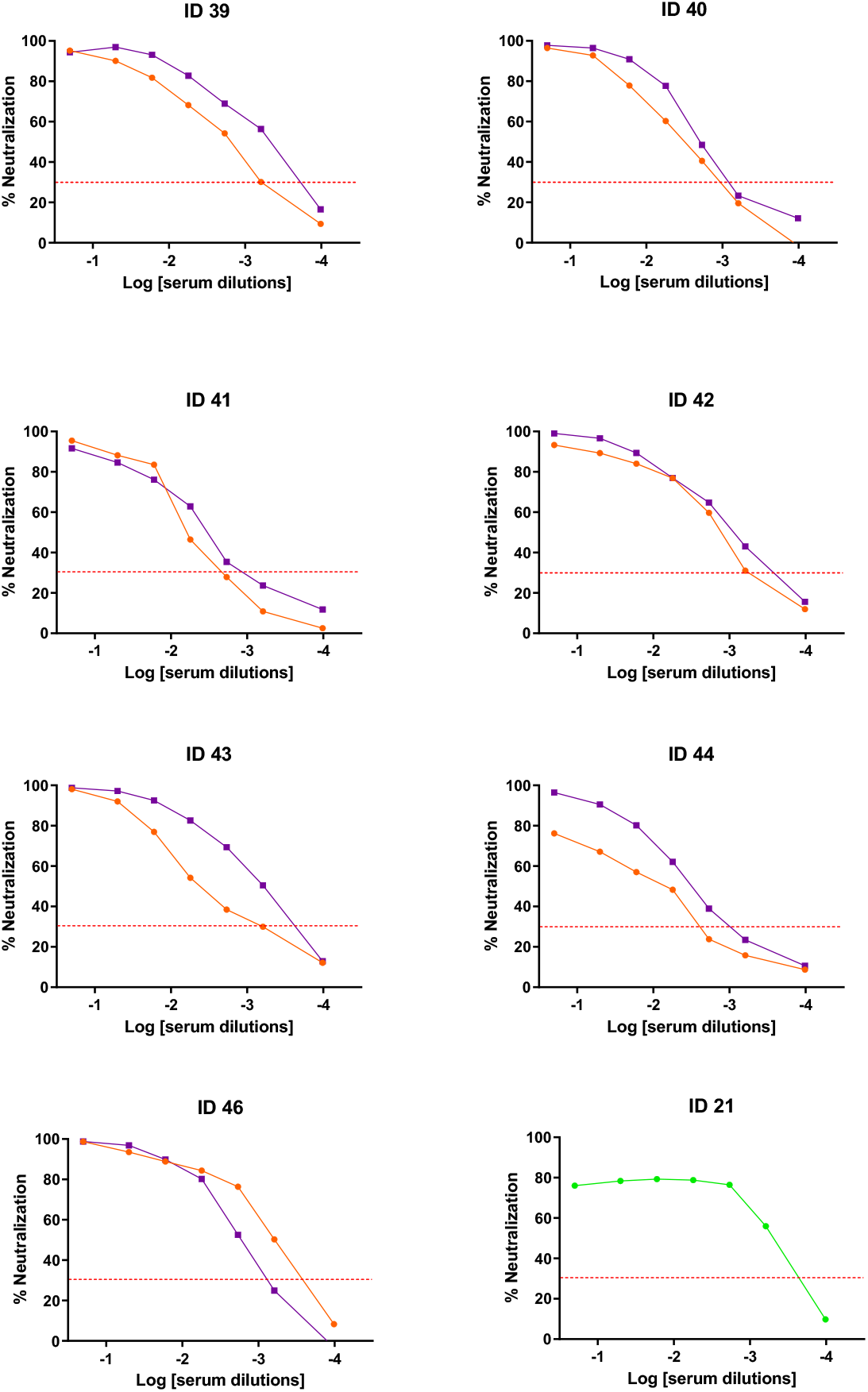
Example of individual patterns of neutralization. Orange: first dose of BNT162b2 vaccine, purple: second dose BNT162b2 vaccine, green: COVID-19 sera sample. Dotted line represents an arbitrary 30% neutralization cutoff.

### Comparison between NAb titers and anti-SARS-CoV-2 S IgG concentrations

To compare SARS-CoV-2 NAb and total IgG titers, 14 sera from either vaccinated (Pfizer-BioNTech, 2^nd^ dose) or COVID-19 diagnosed individuals were selected to measure total SARS-CoV-2 S IgG from samples that together exhibited a spectrum of neutralization ranging between 58% and 97.5%. Total IgG against SARS-CoV-2 S1+S2 were measured in the aforementioned samples, and in 7 randomly selected pre-pandemic serum samples, using a quantitative ELISA. Vaccinated and COVID-19 samples contained similar average titers of anti-SARS-CoV-2 S-specific IgG antibodies (Fig. 7A, 0.095 and 0.085 mg/L, respectively). These concentrations were on average 30% higher compared to titers measured in pre-pandemic sera (Fig. 7A). A similar increase over naïve serum concentration has been previously observed using an ELISA specific for SARS-CoV-2 S1^40^. A significant difference in total IgG concentration was evidenced between the vaccinated group and pre-pandemic group only, as the COVID-19 group presented a larger distribution of concentrations. Larger differences have been described in anti-SARS-CoV-2 S total IgG between pre-pandemic sera and sera from individuals exposed to SARS-CoV-2 antigens^41^. To investigate neutralization in these samples, we used the pseudotyped VP-based neutralization assay developed here. As evidenced earlier, an overall significantly higher neutralization was observed for both COVID-19 and vaccinated groups compared to the pre-pandemic group (p = 0.0006 for both comparisons). In addition, there was significantly more neutralization from vaccinated samples compared to individuals exposed to the virus through infection (Fig. 7B, p = 0.0006), in contrast with total IgG concentrations between these groups. Higher titers of anti-SARS-CoV-2 NAb in individuals vaccinated with BNT162b2 compared to COVID-19 patients have been extensively described^16,42^. In summary, after exposure to the antigen, a wide range of concentrations of anti-SARS-CoV-2 S total IgG could be measured while neutralization was restricted between 58% and 97.5% (Fig. 7C). Others have similarly evidenced a range of concentrations for total IgG between 10-100 mg/ml for COVID-19 patients while neutralization was constrained to >95%^42^.

**Fig. 7.**
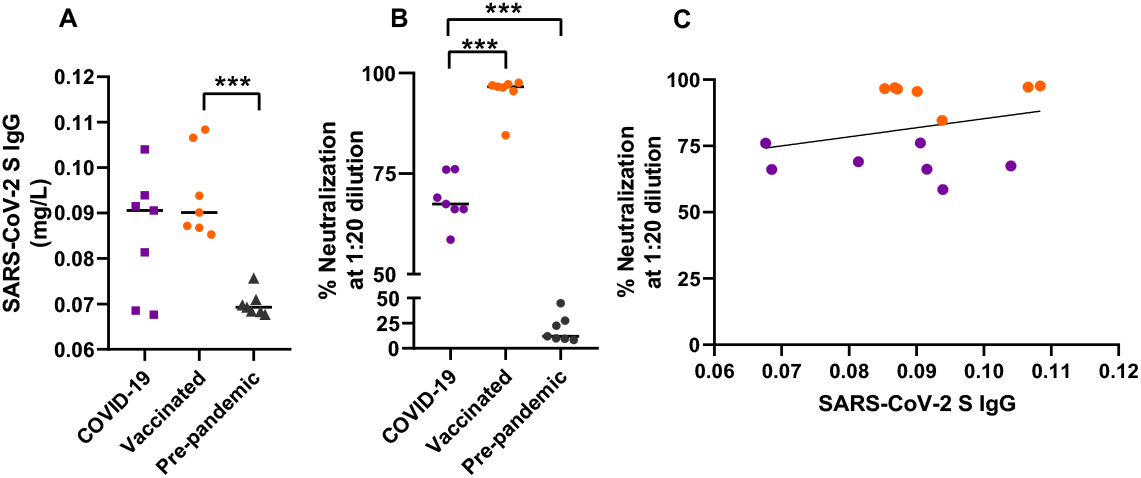
Anti-SARS-CoV-2 S total IgG underestimates neutralization potential evaluated using the SARS-CoV-2 pseudotyped VP-based assay. A: SARS-CoV-2 S ELISA-based quantification of total IgG in COVID-19 convalescent sera, BNT162b2 vaccinated sera and pre-pandemic sera. B: SARS-CoV-2 S VP neutralization by matched sera (used in A) at 1:20 dilution. C: Correlation 1:20 sera neutralization potential vs. SARS-CoV-2 S total IgG. Purple: COVID-19 convalescent sera, orange: vaccinated sera, p<0.001 (***).

## Discussion

As SARS-CoV-2 becomes endemic in human populations worldwide, various selective pressures drive the emergence of viral variants with distinct transmissibility profiles^44^. These variants in turn shape humoral immunity through specific B cell clone selection, which may compromise the efficacy of existing vaccines and increase threshold for herd immunity^45^. In addition, the half-life of SARS-CoV-2 humoral immunity, including NAbs, decays over a few months, regardless of the immunization route (natural infection or vaccination)^6,46^. Mexico has a high prevalence of comorbidities known to increase COVID-19 severity, such as obesity, diabetes and cardiovascular disease, and has experienced higher fatality rates than the global average ^47–49^. Scarce reports are available about SARS-CoV-2 NAb in the Mexican population, with all available data relying on a neutralization-surrogate ELISA kit^15–17^. Herein we propose a BSL-2 safe functional assay to investigate effective humoral immunity against SARS-CoV-2 locally. We produced lentiviral particles bearing SARS-CoV-2 S and optimized a highly sensitive and accurate assay of pseudotyped VP neutralization that can be deployed in most research laboratories in the country to support studies of SARS-CoV-2 induced humoral immunity.

Various pseudotyped VP-based systems have been used worldwide to assess sera- and therapeutic antibody-mediated SARS-CoV-2 neutralization, each with intrinsic protocolar and technical characteristics that have been discussed elsewhere^14,20^. Various assay parameters can be customized, affecting the results. For instance, the neutralization assay presented here has a turn-around time of 24 h after VP inoculation, while others have investigated neutralization as early as 12 h and up to 72 h^44,50^. Longer incubation times increase infection probability therefore comparing neutralization titers obtained using different incubation lengths could be biased^35,51,52^.

Luminescence (as a transduction surrogate marker) increases with the amount of VP applied to target cells. Accordingly, an additional factor affecting results is the differential incorporation of SARS-CoV-2 S on VP membranes, depending on the VP production technique used^23^. Using a SARS-CoV-2 sequence lacking the last 19 amino acids at the C-terminal is known to increase VP membrane incorporations^3^. Therefore, with all other parameters equal, a pseudoviral system using the authentic SARS-CoV-2 S sequence may result in lower RLUs compared to using the SARS-CoV-2 S Δ19 sequence, precluding adequate comparisons.

In this work, we used Nluc as a reporter gene with luminescence as assay read-out. This engineered and enhanced form of luciferase, provides a very sensitive assay, with reports of single cell infections detected^20^. However, furimazine is an expensive substrate. As an alternative, others have reported the development of systems relying on fluorescence measurement to assess transduction^20,53^. These assays may not be as sensitive as luminescence-based methods but could be cheaper when applied in high-throughput screening.

The MOI, or amount of VP added per target cell, is also critical for reaching TCID50 within the timeframe of the assay. While some articles report volumes and dilutions of untitrated viral stock added per well, others titrate VP concentrations to provide a precise MOI^44,54^.

Target cells can be attached to wells at a pre-defined concentration at the time of adding the serum-VP mixture, or alternatively single cell suspensions of target cells may be added to the co-incubated serum-VP mixture^50^. As proteolytic cleavage of ACE2 by ADAM17 and TMPRSS2 affects susceptibility to infection, it is possible that recent trypsin treatment also impact ACE2 cleavage on target cells^55,56^. In this sense it would be relevant to compare assays with similar protocolar details for inoculation (adding VP to adherent cells, or adding freshly trypsinized cells to VP).

Finally, the neutralization threshold used to analyze results is arbitrary, with reports showing analyses using threasholds ranging between 20% and 50% neutralization^23,51,57^. This threshold is used to determine “positivity” of neutralization, and therefore affects reported neutralization titers (last dilution before neutralization curves cross the threshold), and calculated sensitivity and specificity of each assay. The aforementioned pitfalls have highlighted the need for a standardized assay to compare neutralization results across cohorts and worldwide^58^.

We evidenced more variability in the magnitude of COVID-19 neutralization curves, compared to vaccinated sera. Potent NAb clones have been isolated from both high and low neutralizing titers in COVID-19 patients, which suggests SARS-CoV-2 infection hampers appropriate B cell maturation and expansion^59,60^. Much remains to be clarified about the significance of NAb titers. For instance, in COVID-19 patients, NAb titers positively correlate with disease severity^41,57,61,62^. On the other hand, a study reported that about 30% of individuals recovered from mild COVID-19 did not present NAb titers, hinting that other components of the immune system strongly contribute to recovery^59^. As evaluating T cell-mediated immunity to SARS-CoV-2 *in vitro* remains a challenge, monitoring NAb is mandatory to provide clues needed to elucidate immune requirements for protection against, and recovery from, SARS-CoV-2 infections. Vaccine development also requires an easily adapted and safe-to-use platform to measure the induction of immune response, in particular, the detection of NAbs generated in response to the inoculated antigen. The assay developed in this work could be easily adapted to emerging variants by either applying directed mutagenesis to the S sequence, or replacing it with a synthetic gene^53^. As vaccines may need to be adapted to target emerging variants of concern, effective immunity brought by novel vaccines and variant-mediated infections can be monitored locally using this assay. We foresee the proposed platform will shed light onto the development and characterization of humoral immunity to SARS-CoV-2 in Mexico and Latin America.

## Material and Methods

### Vector constructions

The pCAG-HIVgp and pCMV-VSV-G-RSV-REV plasmids were acquired through the Riken Institute BioResource Center^53^. The pCMV-SARS-CoV-2-S-RSV-REV plasmid was produced by cloning the SARS-CoV-2 S sequence, obtained from pCMV14-3X-Flag-SARS-CoV-2 S which encodes codon optimized SARS-CoV-2 S protein lacking the last 19 amino acids at the C-terminal^3^, in place of the VSV-G gene, between the *Nhe*I and *Xba*I restriction sites. The pLenti-Nluc was produced by cloning the Nluc sequence amplified from pCCI-SP6-ZIKV-Nluc, between the *Xba*I and *Bam*HI restriction sites within the pLentiCRISPR v2 backbone (cat. 52961, Addgene, Massachussets, USA.) and removing the Cas9 gene. We produced the plasmid pCMV-REV by removing the envelope protein gene from pCMV-VSV-G-RSV-REV. All constructions were verified by restriction mapping and validated by Sanger sequencing (Sup. Fig. 1).

### Cell culture

HEK-293T (ATCC CRL-3216), Vero (ATCC CCL-81), Vero E6 (ATCC CRL-1586) and Caco-2 (HTB-37) cells were obtained from ATCC and maintained in high-glucose DMEM (Caisson, cat. DML10) supplemented with 10% heat inactivated FBS (Sigma, Missouri, USA. cat. F2442) at 37 °C in a 5% CO_2_ atmosphere. Cells were passaged according to provider’s instructions, using either gentle scrapping or brief exposure to trypsin (Hyclone, Massachussetts, USA. cat. C838R55). All cell lines were used before passage 25.

### Production of SARS-CoV-2 Spike-expressing lentiviral particles

Plasmids pLenti-Nluc, pCAG-HIVgp, and pCMV-SARS-CoV-2-S-RSV-REV were co-transfected at a 3:2:1 DNA ratio using the calcium phosphate method to confluent HEK-293T cells. This transfection protocol was also used to generate the other VP used in this work (without glycoprotein or with VSV glycoprotein). Transfected cells were incubated at 37 °C with 5% CO_2_ for 24 h before replacing the medium to DMEM with 10% FBS. VP-containing supernatant was collected at 72 h post-transfection, clarified by centrifugation, filtered through a 0.45 μm filter (GE Healthcare, cat. 67802504), aliquoted and stored at -80 °C until use. For each production batch, one aliquot was titrated after a single freeze-thaw cycle using the QuickTiter Lentivirus Titer Kit (Cell Biolabs, California, USA. cat. VPK-107) according to the manufacturer’s protocol.

### Flow cytometry

ACE2 expression was measured on all cell lines by flow cytometry using a FACS Celesta fitted with 405 nm, 488 nm and 533 nm lasers (BD Biosciences). Briefly 3 x10^5^ cells were incubated on ice with mouse anti-human ACE2 monoclonal antibody conjugated to AF-647 (Santa Cruz Biotechnology, USA, cat. SC-390851) following manufacturer’s instructions. Propidium iodide (BD Biosciences, cat. 51-6621E) was added 15 min before acquisition on the flow cytometer, as per manufacturer’s instructions. At least 20,000 events were acquired per sample, and the data was analysed using FlowJo v.10.

### Transduction assay

Transduction of target cells by VP was assessed by measuring the level of luminescence induced by the conversion of furimazine, reported in RLUs, using a commercial kit according to manufacturer’s instructions (Promega, USA. cat. N1110), including reading at 460 nm on a Biotek Synergy Microplate Reader. Uninfected cells were used for normalization.

### Western blot

The selective incorporation of SARS-CoV-2-S or VSV-G, and p24 proteins in VP was validated by western blot. Briefly, supernatants were pelleted by ultra-centrifugation at 25,000 g for 2 h over a 20% sucrose cushion. The supernatant was removed, and the viral pellet was resuspended in 60 μl of PBS. Thirty μl of each sample were subjected to SDS-PAGE followed by immunoblotting on PVDF membranes. Mouse anti-VSV-G-HRP (Santa Cruz, cat. SC-365019-HRP) and mouse anti-HIV1-p24-HRP (Santa Cruz, cat. 69728-HRP) were used for one-step detection. A chimeric monoclonal antibody (Sino Biological, cat 40150-D001) was used to detected SARS-CoV-2-S as primary antibody, and a goat anti-human IgG conjugated to HRP (Bio-Rad, cat. 204005) was used as secondary antibody. Membranes were revealed using the Immobilon Forte Western HRP substrate (Millipore, cat. WBLUF0500) following the manufacturer’s protocol. Chemiluminiscence signal was acquired through ChemiDoc XR S+ (Bio-Rad).

### TCID50 determination

TCID50 assays were performed as previously described^15^. Viral stocks were serially diluted 3-fold and each dilution assessed in 6 replicates in the infection of 25,000 Vero cells per well, seeded in a 96-well plate and incubated at 37 °C in a 5% CO_2_ atmosphere. Twenty-four h post-seeding, luminescence was measured as a surrogate for infection as described above.

### Pseudovirus-based SARS-CoV-2 Spike neutralization assay

Protocols for the use of human samples for this work were approved by the IRB of the Instituto Mexicano del Seguro Social (IMSS) with reference number R-2020-785-068 prior to starting this work. A total of 15 SARS-CoV-2 free, collected between 2014 and 2018 (prior to the 2019 initial outbreak, collected through IRB approved protocols at IMSS, for bio banking purposes), and 18 COVID-19 blood samples were obtained from 15 patients (confirmed by a positive qRT-PCR diagnostic, Tables 1). Samples were collected and used upon signed informed consent and anonymization. Vaccinated sera were obtained from health care professionals receiving the BNT162b2 vaccine. Blood samples were collected between 14 and 16 days post-application of the first dose, and a second sampling was performed 27 to 29 days after the application of the second dose. Status of prior infection with SARS-CoV-2 was also recorded (Table 2). Briefly, sera were enriched from coagulated blood by centrifugation, inactivated for 30 min at 56 °C, and aliquoted and stored at -80 °C until use. On the day of the assay, sera were serially diluted 7-folds, spanning 1:5 to 1:9860, and 100 μL of each dilution was incubated for 1 h at 37 °C and 5% CO_2_, together with 15 pg of SARS-CoV-2 VP in duplicate in a 96-well plate. Post-incubation, 25,000 Vero cells were added to each well and the plate was incubated for 24 h at 37 °C and 5% CO_2_. As a positive control for transduction, SARS-CoV-2 VP were incubated with Vero cells. Vero cells seeded in triplicate were used for basal luminescence background assessment as described earlier. After 24 h, Nluc levels were measured as described above. Neutralization is described as % inhibition of transduction, calculated as: Inhibition (%)=(mean RLUs of infected control wells – mean RLUs (duplicate) of sera-treated well) x100.

### Statistical analysis

All statistical analyzes were performed using GraphPad Prism v.9 software and p-values < 0.05 were considered statistically significant. In the case of categorical values for calculation of Sn and Sp, a contingency 2×2 table was used with a fisher exact test. For flow cytometry analyzes, medians of fluorescence intensity and percentage of positive cells were compared using Mann-Whitney and P values are reported. To estimate EC50, neutralization curves were log transformed, normalized, and fitted to the most appropriate model between log(inhibitor) vs. response (three parameters) and log(inhibitor) vs. response, variable slope (four parameters). For calculating sensitivity and specificity of the assay, we determined sera positivity and negativity at a final 1:20 dilution and using a 30% neutralization threshold, as previously reported to evidence true/false positives and true/false negatives^43^.

## Acknowledgements

We warmly thank Dr. Zhaohui Qian (Institute of Pathogen Biology, Beijing, China) for sharing pCMV14-3X-Flag-SARS-CoV-2 S with us. We acknowledge the financial support received by CONACYT (scholarships 1007842 and 657487), Secretaría de Relaciones Exteriores, Tecnológico de Monterrey and StrainBiotech S.A de C.V, that made this work possible.

---

**Sup. Fig. 1:**
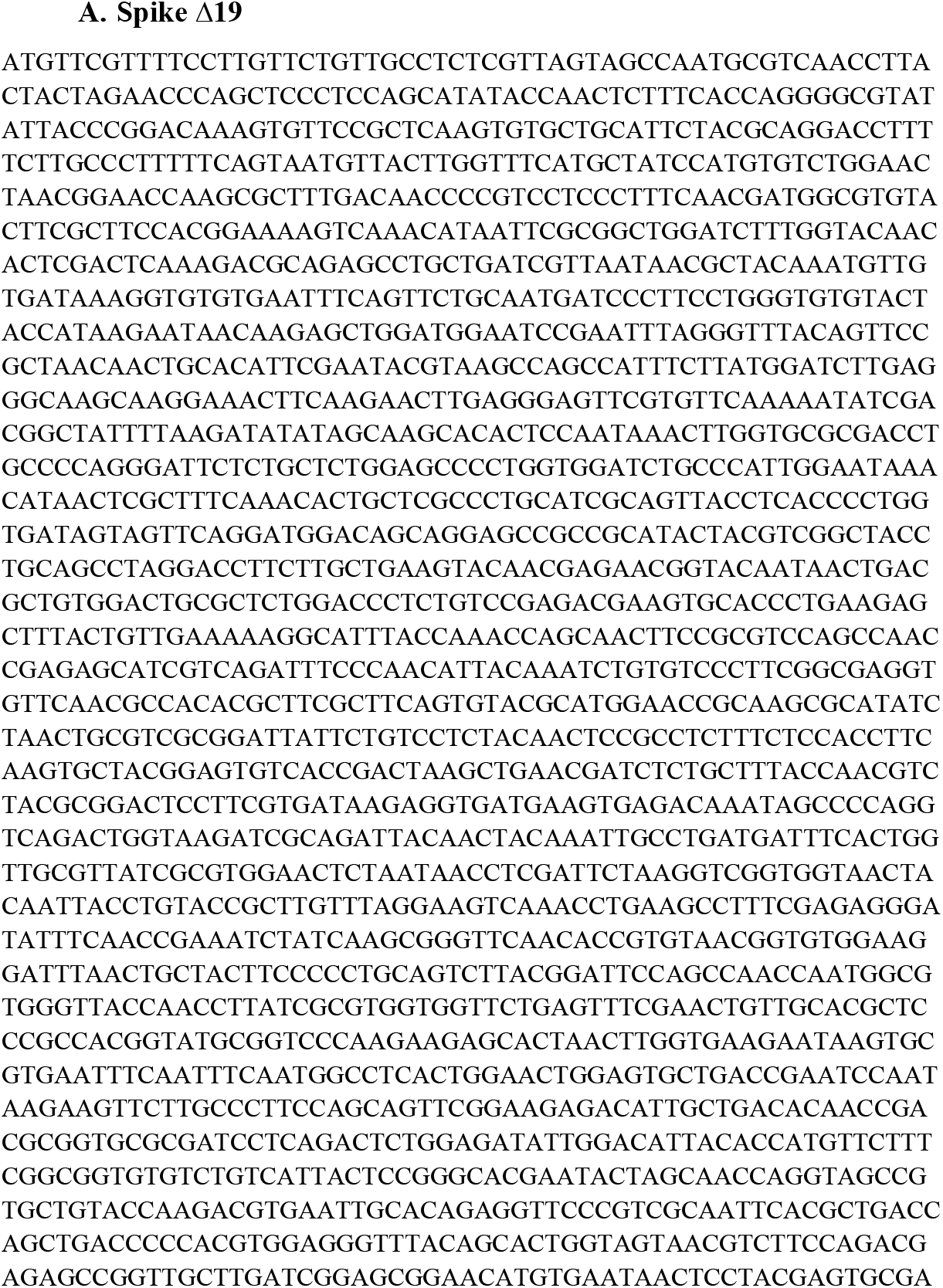
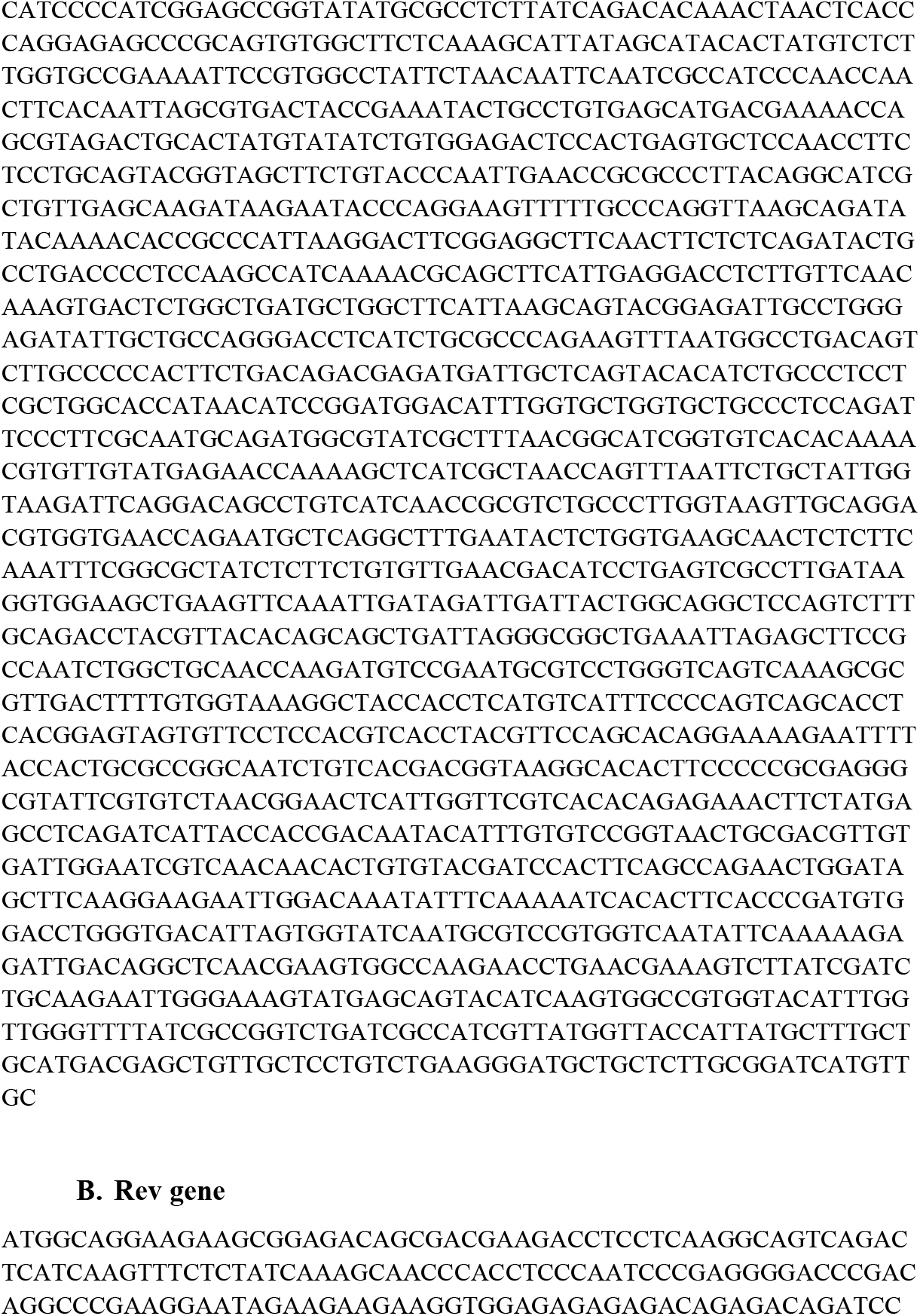
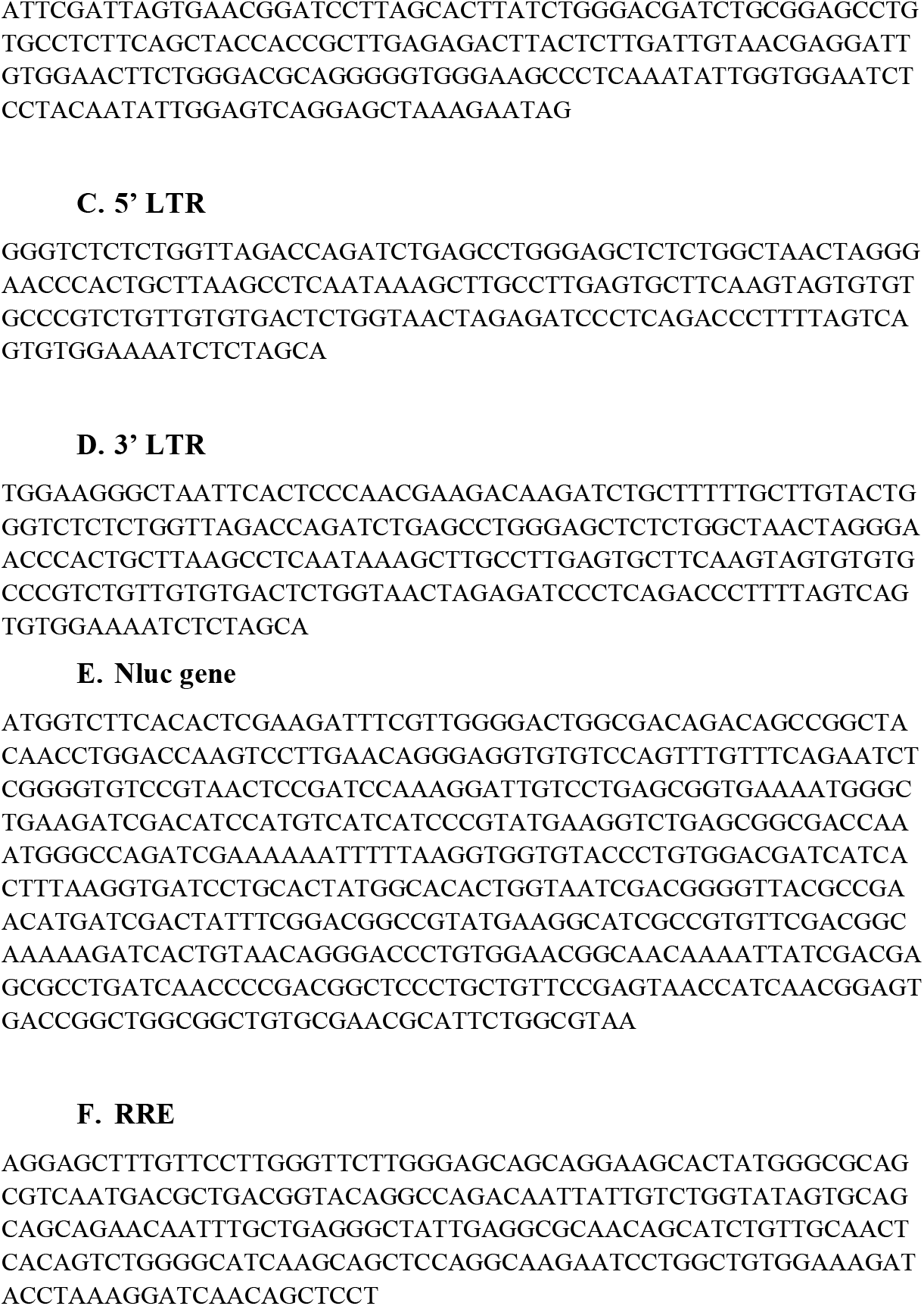
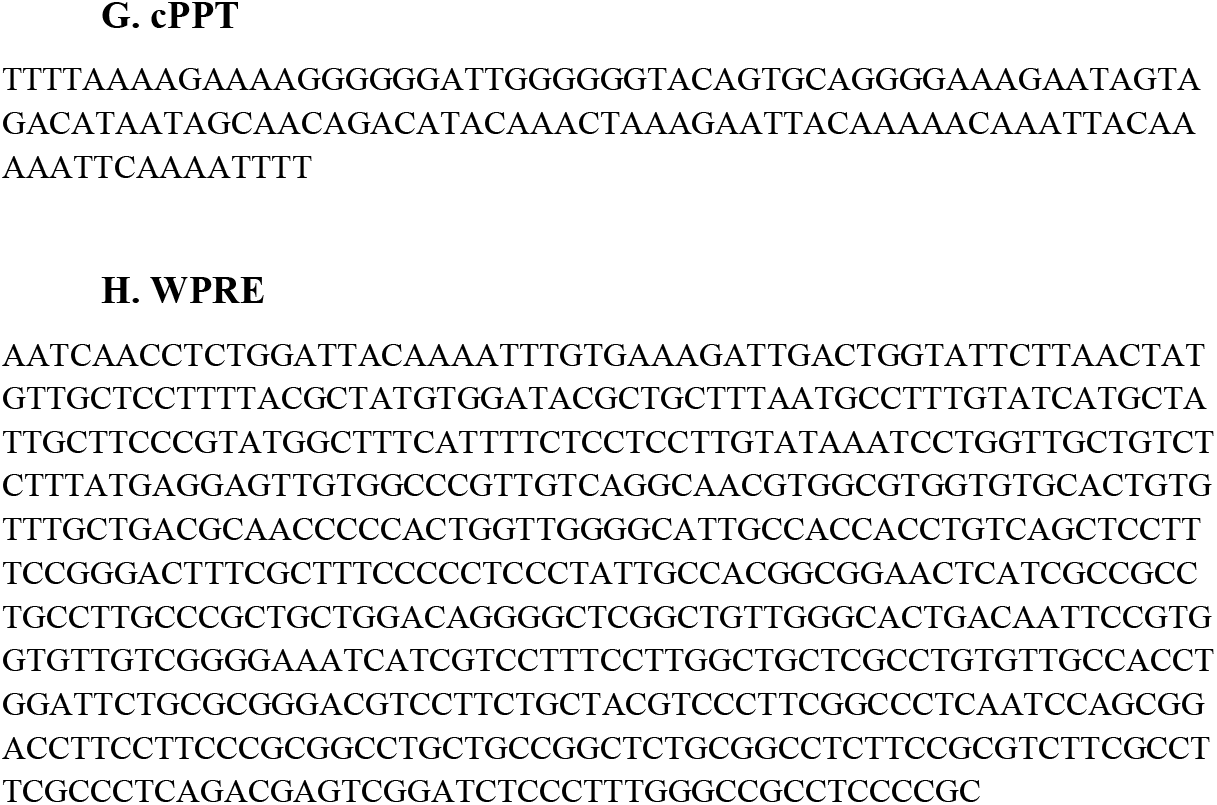
Sequences of relevant plasmid portions obtained through Sanger sequencing

**Sup. Fig. 2:**
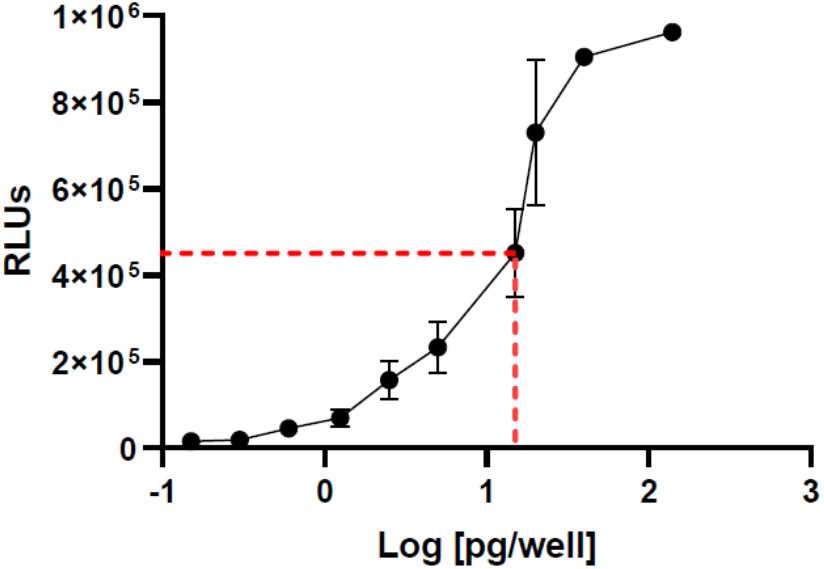
Determination of the TCID_50_ of SARS-CoV-2 S pseudovirus using 25,000 Vero cells as target in 96 well plate with infection assessed at 24 h. The dotted line represents 50% of maximal RLU equivalent to 15 pg VP per well. The average of 2 independent experiments, ran in triplicate, is shown. Error bars indicate SD

